# A comprehensive quality control pipeline in human microbiome research for large population studies

**DOI:** 10.1101/2025.01.20.633643

**Authors:** Ruolin Li, Joost A.M. Verlouw, Cindy G. Boer, Pascal Arp, Joyce van Meurs, Andre G. Uitterlinden, Robert Kraaij, Carolina Medina-Gomez

## Abstract

The widespread application of high-throughput Next Generation Sequencing (NGS) technologies has made microbiome research an emerging field in public health and biomedical sciences. However, there are still many challenges that need to be addressed in this field. Pipelines available to generate microbiome data across cohorts are diverse, and sources of variation to be recorded and evaluated during microbiome profiling have not been standardized. Moreover, meticulous quality control of the microbiome data processing, from collection to computational quantification is still challenging, especially in large population studies. Innovative approaches are required to handle samples and to minimize the potential bias introduced by logistic hurdles in biobanking. In this paper, we describe the methodological steps surrounding the optimization of the 16S rRNA gut microbiome profiling in two large prospective cohorts the Generation R Study (mean age 9.83 ± 0.32 years) and the Rotterdam Study (mean age 62.67 ± 5.66 years). This paper also highlights potential solutions to sample mislabeling in large-scale microbiome analysis. To summarize, our study addresses common problems in human microbiome research. It aims to improve the research quality and reliability by integrating more stringent quality control standards into microbiome research.

## 1 Introduction

The human body is colonized by a vast number of bacteria, archaea, microbial eukaryotes and viruses that are as abundant as our somatic cells, of particular interest are these communities harbored in the gut. The gut microbiome can impact human physiology through different biological processes. Due to its high diversity in composition and function, the gut microbiome greatly expands our capacity to metabolize food and drugs [1, 2], as well as to shield against xenobiotic toxins [3]. In line with these facts, a large body of evidence now supports the role of the gut microbiome in our predisposition for various diseases including obesity [4], type 2 diabetes mellitus (T2D) [5], hypertension [6], hyperlipidemia [7], cardiovascular disease (CVD) [8], osteoporosis and osteopenia [9], sarcopenia [10], neurodegenerative diseases [11], and depression [12]. Therefore, the gut microbiome has become a crucial matter, generating growing interest from the clinical and biomedical research community.

A widely used method for characterizing bacterial communities is high-throughput sequencing of the hypervariable regions of the 16S rRNA gene. Even if metagenomics sequencing technology is available and offers greater precision, amplicon-based 16S rRNA sequencing is still broadly applied in large-scale population studies [13–15] due to its lower cost. The development of online platforms such as MicrobiomeDB [16] has further facilitated the integration and comparison of data across multiple microbiome studies. Despite its widespread adoption over the past decade, the application of 16S sequencing in epidemiologic research continues to face several challenges, as previously summarized [17]. Sources of variation that can influence microbiome-health association analyses may arise at every stage of the workflow—from fecal sample collection and storage, to DNA extraction, PCR amplification of the 16S rRNA gene, sequencing, bioinformatic processing, and statistical analysis. Therefore, rigorous documentation and standardization of experimental procedures are essential to ensure reproducibility and reliability in microbiome research.

Seven years ago, we published the first profiling of the stool microbiome from children and adults participants of two large, independent, and well-characterized population-based cohorts [18]: the Generation R Study (GenR) [19] and the Rotterdam Study [20] (RS, sub-cohort RSIII). These datasets have been widely used to explore associations between the gut microbiome and various health outcomes, including circulating metabolites [21], depressive symptoms [22], T2D [23], hepatic steatosis [24], or joint pain and inflammation [25] in adults, and atopic diseases [26] in children. They have also supported investigations into the long-term effects of antimicrobial drug use [27] and the influence of host genetic factors on gut microbiota composition [28].

The need to update this initial release arose from advancements in amplicon sequence processing, particularly the introduction of denoising algorithms such as Deblur [29] or Divisive Amplicon Denoising Algorithm pipeline (DADA2) [30], which improve taxonomic resolution by reducing false positives. Additionally, updated reference taxonomies like SILVA 138.1 [31] or GreenGenes2 [32], have enhanced classification accuracy. These developments provided an opportunity to revise and improve our microbiome analysis pipeline. Furthermore, we developed innovative quality control (QC) procedures to ensure more reliable and traceable research datasets. In particular, we addressed the challenge of sample swaps—a common issue in biomedical research [33] that can weaken associations, reduce statistical power, or introduce bias in epidemiological studies [34]. In this paper, we present the updated GenR and RS 16S rRNA sequencing microbiome datasets, finalized in July 2021. We detail the experimental methodology, beginning with DNA-isolated samples previously stored, and describe the steps taken to validate and profile the microbiome data.

## 2 Methods

### 2.1 Description of the study populations

The Generation R Study is a population-based prospective multi-ethnic birth cohort study, conducted in Rotterdam, the Netherlands, designed to identify early environmental and genetic factors underlying normal and abnormal growth, development, and health before adulthood [19]. Data on mothers and their children were collected through multiple follow-up rounds. Stool sample collection began in 2012, when participating children had a mean age of 9.8 years (SD: 0.3) [18]. A total of 2,921 samples were received at the Genomics Core Facility (CoFa) of Erasmus Medical Center Rotterdam (Erasmus MC) for microbiome profiling. The study received ethics approval from the Medical Ethical Committee of Erasmus MC (MEC-2012-165), and written informed consent was obtained from all participants’ parents. All procedures were conducted in accordance with the Declaration of Helsinki.

The Rotterdam Study is a prospective population-based cohort initiated in 1990 to study determinants of disease and disability in Dutch adult individuals [20]. RS consists of four sub-cohorts and includes approximately 18,000 residents of the Ommoord suburb in Rotterdam, primarily of European ancestry, aged ≥ 40 years. The collection of fecal samples started in 2012 among the RS-III sub-cohort at a mean age of 62.7 years (SD:5.7 years) [18]. In total 1,691 samples were received in the CoFa facility of Erasmus MC for profiling. All subjects provided written consent prior to participation in the study. All protocols were performed in accordance with the Declaration of Helsinki. Ethics approval was obtained from the Medical Ethical Committee of Erasmus MC (MEC-02-1015).

### 2.2 Previous release of the Rotterdam Study and Generation R Study microbiome data

The protocol used to generate the microbiome datasets for the Generation R Study and the Rotterdam Study—from sample collection to taxonomic profiling—has been previously described in detail [18]. Briefly, stool samples were collected at home by the participants using a Commode Specimen Collection System (Covidien, Mansfield, MA) and sent through regular mail to the lab at the Erasmus Medical Center. Upon arrival, samples were immediately stored at −20 °C. DNA extraction was performed using the Arrow Stool DNA kit (DiaSorin S.P.A., Saluggia, Italy) on homogenized, bead-beaten samples [18]. DNA concentrations were quantified using Quant-iT PicoGreen dsDNA Assay Kit (Thermo Fisher Scientific, Waltham, MA). DNA was stocked in 96-well plates in which 4-wells contained only purified water used as controls. Next, the V3 and V4 variable regions of the 16 S rRNA gene were amplified, normalized using the SequalPrep Normalization Plate kit (Thermo Fischer Scientific), and pooled. Amplicon pools were purified using the Agencourt AMPure XP (Beckman Coulter Life Science, Indianapolis, IN), and their size and concentration were assessed using the LabChip GX (PerkinElmer Inc., Groningen, The Netherlands). Sequencing was performed on an Illumina MiSeq platform (Illumina Inc., San Diego, CA, USA). Phylogenetic multi-sample profiling was performed on QIIME (version 1.9.0) and UPARSE (version 8.1) software packages using SILVA rRNA version 128 as the reference [18]. Fastq files were subsampled to 10k reads using an in-house script. Samples that were below the threshold of 10k reads were excluded. This pipeline was based on the construction of molecular operational taxonomic units (OTUs). After applying a strict quality control (QC), the final dataset included microbiome profiles from 2,111 participants in the GenR and 1,427 participants in the RS [18].

### 2.3 Generation of the new microbiome data release

#### 2.3.1 Processing of raw sequencing reads

We reprocessed the previously generated raw sequencing datasets—prior to QC—using the Divisive Amplicon Denoising Algorithm implemented in the DADA2 R package (version 1.18.0) [35]. Initially, barcodes were removed from the reads, and paired-end reads were concatenated. These barcodes were then used to demultiplex the reads, allowing a maximum of one mismatch per 12-nucleotide segment (half of the 24-nucleotide barcode). Primer sequences and heterogeneity spacers were removed using TagCleaner (version 0.16) [36]. Trimmed reads were imported into the DADA2, and samples with no remaining reads were excluded. The DADA2 quality filter removed reads with an expected error rate higher than two in either the forward and reverse reads, as well as reads containing ambiguous bases. Additionally, reads were truncated at the first occurrence of a low-quality base (Q-score =<2). Denoising was performed by clustering reads based on sequence similarity, beginning with the most abundant read. Reads within 10% similarity were aligned to the cluster and retained if their abundance p-value (i.e., indicating the likelihood of being erroneously abundant) was below the default threshold. This process was repeated with the iteratively for all remaining reads. Singleton reads were automatically excluded due to the algorithm’s requirement for multiple identical sequences. Forward and reverse reads were merged only if they overlapped with a 100% match; otherwise, they were discarded. Reads truncated during initial filtering were often removed at this stage due to insufficient overlap. The resulting merged reads were used to construct an amplicon sequence variant (ASV) table. Chimeric ASVs were identified and removed using the removeBimeraDenovo function in consensus mode within DADA2. After applying this process to further refine the dataset, we applied abundance and prevalence filters: ASVs were retained only if they represented at least 0.0005% of the total cleaned reads and were present in at least 1% of the samples.

#### 2.3.2 Identification of sample swaps

The update of the microbiome profiling pipeline at our facility also provided an opportunity to revisit the entire experimental workflow. As part of this revision, we aimed to identify and exclude potential sample swaps and to recover previously excluded low-quality samples.

##### • Swap identification based on host genetic markers

The first step of our new pipeline was to dilute the DNA stocked in plates, as outlined in Section 2.2, to a concentration of 20 ng/µl. To determine the sex of the host, we used the Investigator Quantiplex Pro kit (Qiagen) [37], a real-time PCR assay that quantifies both total human and male-specific DNA. For each sample, 40 ng of DNA was amplified using 384-well PCR plates on the QuantStudio 7 system (ThermoFisher). Samples with less than 2 pg of autosomal DNA were excluded to ensure data quality. A standard curve was constructed to quantify both autosomal and Y-chromosome DNA, enabling accurate sex determination. The genetically inferred sex was then compared to the sex reported in participant questionnaires. Discrepancies were flagged as potential sex-discordant sample swaps.

Because sex-discordant swaps only account for half of all possible sample swaps, we implemented additional strategies to detect same-sex swaps. First, we leveraged genotype data from blood samples of RS and GenR participants for host identification. To assess feasibility, we selected the 15 Generation R samples with the highest host DNA concentration—quantified using the Investigator Quantiplex Pro kit—and genotyped them using the Illumina Global Screening Array v1 (GSA v1). We then performed a concordance analysis using 36,629 biallelic single nucleotide polymorphisms (SNPs) shared across the HapMap reference panel [38], the GSAv1 array and the Illumina 610 array, which had previously been used to genotype blood-derived samples from GenR participants.

In parallel, we identified samples from both cohorts with available genotype data and stool-derived autosomal DNA concentrations exceeding 90 pg. For these samples, we developed a contingency plan to enable individual identification using stool-derived DNA, which was often limited and potentially degraded. Specifically, we selected a panel of 12 TaqMan assays (ThermoFisher)—rs225014, rs1799722, rs1042713, rs6280, rs10948155, rs11755164, rs1990622, rs12462534, rs7597593, rs1537372, rs2228145, rs514659—based on their presence in both RS and GenR cohorts, either through direct genotyping or high-confidence imputation (MACH r² > 0.9). These assays were designed to be run on stool-derived DNA to enable cross-platform genotype concordance checks. To evaluate the reliability of this approach, we compared genotypes obtained via TaqMan assays with genome-wide genotypes from array-based platforms.

##### • Swap identification based on microbiome similarity-distance matrix

To further identify sample swaps, we developed a pilot approach based on microbiome similarity metrics by re-sequencing a random subset of samples from the initial data release [18]. This method was grounded in the assumption that microbiome profiles from the same individual should exhibit the highest similarity within the cohort [39, 40].

We randomly selected 350 samples each cohort collection. Of these, 211 GenR and 150 RS samples had sufficient DNA remaining on the diluted plates and were not previously flagged as sex-discordant swaps. To validate the performance of the algorithm, we also included 15 duplicate pairs from GenR, and 7 from RS. All samples, original and re-sequenced, were processed using the updated pipeline. Samples with fewer than 4,000 reads were excluded, and the remaining samples were rarefied to this threshold.

Beta-diversity was calculated using Aitchison distance on centered log-ratio (CLR) transformed ASV tables, with a pseudo-count of one added to handle zeros. For each re-sequenced sample, we computed its similarity to all other samples within the same cohort and plotted the resulting distance distribution (**Figure 1**). If the re-sequenced sample did not most closely match its original counterpart but instead had outlying proximity to another sample, it was flagged as a potential swap and investigated further. To ensure robust interpretation, we reviewed metadata annotations for flagged samples, including prolonged time between sample deposition and freezing (TimeInMail) and notes on DNA amplification issues. These factors were considered potential artefacts influencing beta-diversity and were accounted for in the evaluation. An illustrative example of this swap identification process is shown in **Figure 1**.

**Figure 1.**
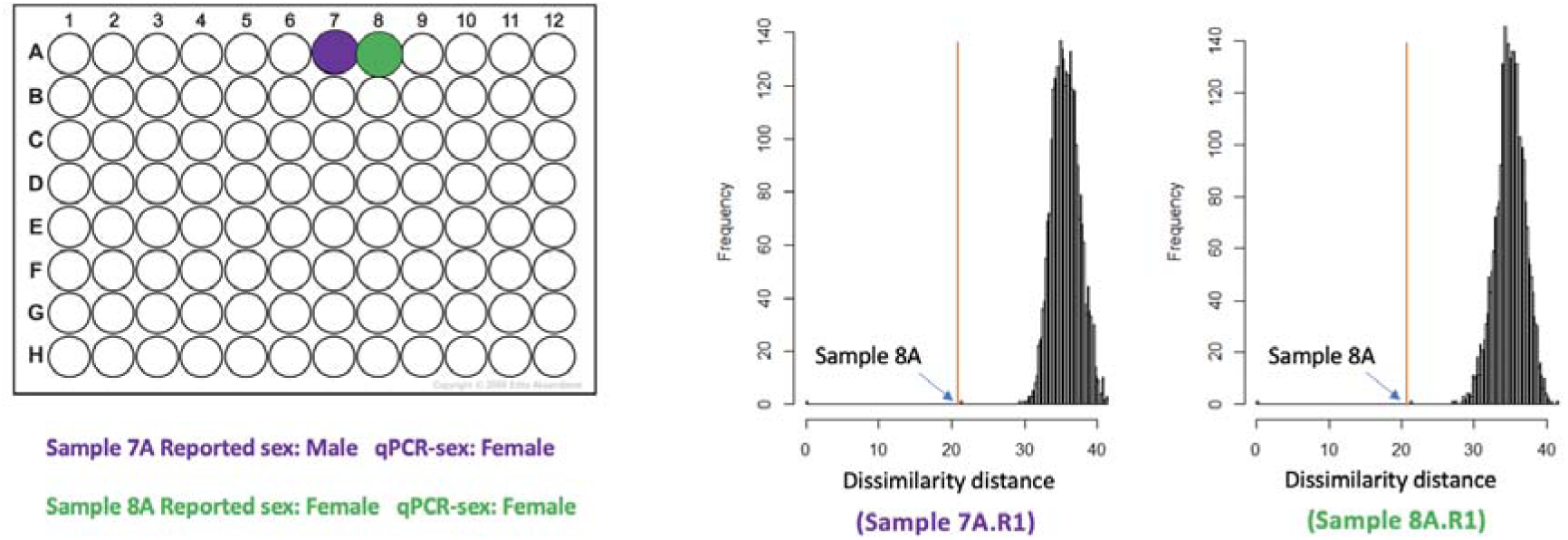
Overview of the swap identification pipeline. Following qPCR for sex determination on host-DNA in one of the 96-well dilution plates, sample 7A was genetically identified as female, in contrast to participant records indicating male sex. The histogram of microbiome distances (right panel) shows that the re-sequenced sample 7A.R1 is most similar to adjacent sample 8A, which was confirmed to be female by both qPCR and questionnaire data. This discrepancy suggests a potential cross-sample swap between 7A and 8A.

Confirmed swaps were defined as cases in which both the original and re-sequenced samples were present in the dataset and did not match. When only one of the two samples was available, potential swaps were identified based on outlying proximity to another sample in the cohort.

Based on insights from the pilot study, we expanded our quality control procedures to further identify and resolve sample swaps. We first identified plates in both cohorts that were enriched for potential swaps and subjected them to a second round of re-sequencing. If the microbiome profiles from the two sequencing rounds did not match (based on similarity distances), the sample was excluded. If samples matched, then the one with highest read count was retained. Next, we reviewed laboratory logs from the previous data release [18], to identify plates with a high number of samples previously excluded during quality control. These plates were prioritized for a third round of sequencing. Sample selection for re-sequencing was guided by several criteria: (1) the possibility of swaps occurring prior to DNA plate preparation, (2) longer collection time (TimeInMail > 7 days), and (3) increased risk of manual pipetting errors, particularly for isolated samples in non-contiguous plate positions (e.g., single interior wells), which pose higher error risk compared to systematic pipetting patterns such as contiguous rows, columns, or edge positions. After processing through the bioinformatics pipeline, samples with more than 4,000 reads were retained; those below this threshold were excluded. Additionally, samples with prolonged TimeInMail— defined as >7 days for RS and >11 days for GenR—were excluded from the final dataset due to potential degradation. An overview of this workflow is presented in **Figure 2a**.

**Figure 2.**
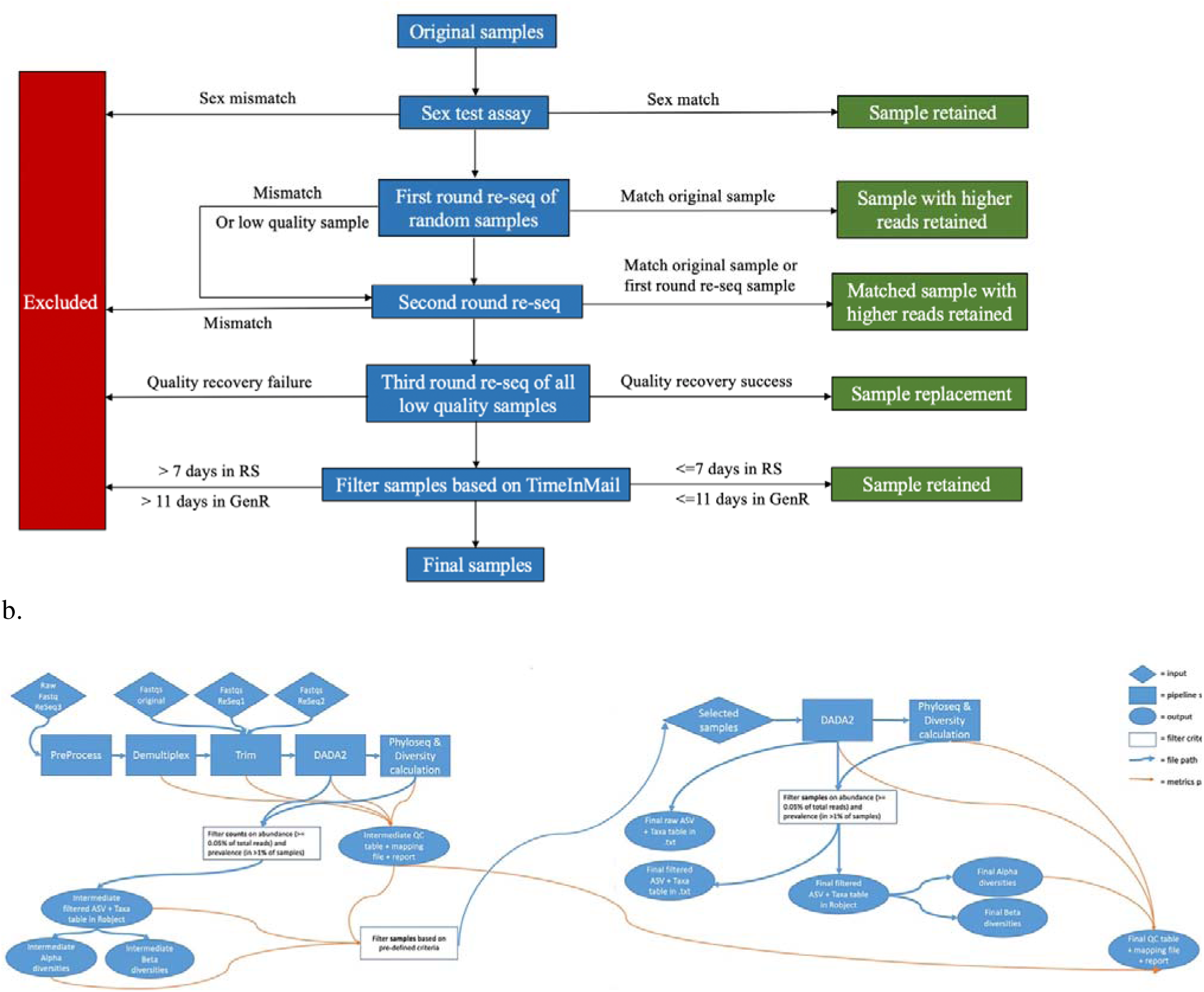
Overview of the Quality control (QC) workflow. (a) Re-sequencing and sample filtering process. This panel illustrates the sequential QC steps applied to the microbiome datasets, including sex determination via qPCR, first-, second-, and third-round re-sequencing, and filtering based on TimeInMail. Workflow steps are shown in blue, retained samples in green, and excluded samples in red. Low-quality samples were defined as those with fewer than 4,000 sequencing reads. **(b) Bioinformatic QC pipeline.** This panel outlines the generation of the updated microbiome datasets for the Generation R Study and the Rotterdam Study. It highlights the sequential sample filtering criteria based on pre-defined criteria: 1) samples with less than 4,500 reads, or 2) samples with 4,500–6,000 reads that lost more than 20% of reads during abundance and prevalence filtering, or 3) samples with >6,000 that lost more than 50% of reads during abundance and prevalence filtering. File generation steps are indicated with blue arrows, and metric flow paths with orange arrows.

#### 2.3.3 Additional Quality Control Steps in the Bioinformatic Pipeline

Beyond ASV-level filtering (abundance and prevalence criteria described above; **Figure 2b**), we implemented sample-level quality filters to identify samples with low sequencing depth or disproportionate loss of reads during ASV filtering, both indicative of technical issues affecting data quality.

Through inspection of read count distributions through the DADA2 pipeline and diversity metrics, we observed that samples falling into specific categories exhibited aberrant microbiome profiles characterized by unusually high beta-diversity distances from cohort distributions and inconsistent profiles among technical replicates. Based on these empirical observations, we excluded samples meeting any of the following criteria: a) Low sequencing depth: <4,500 reads after initial quality filtering, below which diversity estimates become unreliable. b) High loss rate in moderate-depth samples: 4,500–6,000 reads with >20% loss during ASV filtering (abundance and prevalence criteria), indicating a high proportion of non-reproducible features (potential contaminants, sequencing errors). c) High loss rate in adequate-depth samples: >6,000 reads with >50% loss during ASV filtering, suggesting extreme contamination or sparse ASV distributions indicative of technical issues despite adequate sequencing.

Cleaned ASV tables were used for taxonomic classification via the RDP naïve Bayesian classifier [41], trained on the SILVA version 138.1 database [31]. Taxonomic assignments from Kingdom to Genus were accepted if the bootstrap confidence was ≥50% (i.e., calculated as the fraction of bootstrap replicates assigned to a certain taxon). Species-level assignments were accepted only when the ASV matched a single species with 100% bootstrap confidence.

#### 2.3.4 Final microbiome datasets for Generation R Study and the Rotterdam Study

The finalized microbiome datasets for the Generation R Study and the Rotterdam Study include all original and successfully recovered samples, with all identified swaps and low-quality samples excluded. To support robust downstream analyses, detailed metadata were incorporated, including information on read counts post-DADA2 processing, DNA isolation batch [18], sequencing batch (i.e., original, re-sequenced), TimeInMail, season of feces production, and reported use of probiotics and antibiotics. These variables allow researchers to account for potential technical and biological confounders in microbiome analyses.

To visualize the microbial composition of the final datasets, taxonomic profiles were aggregated at the phylum level using the *tax_glom* function from the *phyloseq* package. Average relative abundances of the five major phyla were calculated and displayed using donut plots in R 4.0.

### 2.4 Assessment of technical and biological covariates

To quantify the influence of technical and biological factors on microbiome composition, we performed permutational multivariate analysis of variance (PERMANOVA) using the adonis2 function from the R package vegan (v.2.4–6). We examined both technical factors (TimeInMail, DNA isolation batch, sequencing batch, number of reads, and sampling season) and individual phenotypes (age, sex, BMI, antibiotic use, and ethnicity (GenR only)). The analysis was based on Bray-Curtis distance matrices calculated from ASV relative abundance. We tested both univariate and multivariate models using 999 permutations in individuals with complete data for all these variables (2,252 GenR participants and 1,370 RS participants and defined statistical significance at a false discovery rate (FDR) < 0.05.

To validate our findings, we compared the proportion of variance explained by each covariate to results from the Dutch Microbiome Project (DMP) within Lifelines (LL) [42], a large, three-generational population-based biobank from the northern Netherlands. This cohort includes 8,208 individuals (age range: 8–84 years) with extensive phenotypic and multi-omics data [42]. We visualized the proportion of variance explained by each covariate using bar plots, enabling direct comparison across Generation R, Rotterdam Study, and LL cohorts. Although LL data were derived from metagenomic sequencing rather than 16S rRNA gene sequencing, a previous study analyzing diverse host phenotypes in a subset of this cohort found consistent beta-diversity variance explained patterns between these two sequencing approaches [2], supporting the validity of our cross-cohort comparison.

## 3 Results

### 3.1 Genetic host identification based on stool-recovered DNA

In the Generation R Study, 2,851 out of 2,921 samples (97.6%) successfully underwent sex determination using the TaqMan assay. In the Rotterdam Study, this success rate was 85.5% (1,445 out of 1,691 samples). Discrepancies between genetically inferred and reported sex were observed in approximately 3% of GenR samples (n = 81) and 5% of RS samples (n = 72). In RS, household information was available, and 18 of these mismatches (25%) were traced to individuals of the opposite sex residing in the same household, suggesting potential within-household sample swaps. Mapping these discrepancies to plate layouts revealed that certain plates were enriched for sex swaps in both cohorts and these were flagged for further investigation.

Genotype data from blood samples were available for 1,902 GenR participants (65.1%) and 1,454 RS participants (86.0%) with corresponding stool samples. An initial attempt to use the Illumina GSA array for host identification yielded high-quality genotyping (call rate >95%) in only 2 of 15 GenR samples (13%). One sample showed perfect concordance with blood-derived genotypes (call rate 99%), while the other (call rate 97%) showed only 75% concordance and a high rate of homozygous calls, suggesting possible DNA degradation.

Among the 12 pre-selected TaqMan SNP assays, only rs1990622 met the criteria of high imputation quality (MACH r² > 0.9) and a minor allele frequency >0.30 across global populations. Furthermore, only 246 samples in GenR and 126 samples in RS had sufficient host DNA to attempt this assay. Genotyping success for rs1990622 was 51.2% (n = 126) in GenR and 73.0% (n = 92) in RS. Among successfully genotyped samples, eight mismatches (6.3%) were detected in GenR, three of which had been identified as sex-swaps and three others had failed sex determination. In RS, seven mismatches (7.7%) were identified, five of which were confirmed sex-swaps and one had failed sex determination.

### 3.2 Microbiome-Based Identification of Sample Swaps

In the initial pilot study, after excluding samples that failed pipeline filtering or had fewer than 4,000 reads, 184 re-sequenced samples from the Generation R Study and 134 from the Rotterdam Study remained. Among these, 15 duplicate pairs in GenR and seven in RS were included as internal controls. All duplicate samples correctly identified their corresponding pair as the most similar sample within their respective cohort using the similarity matrix algorithm, validating the approach. However, we identified mismatches where the re-sequenced sample did not most closely resemble its original counterpart. In GenR, 9 out of 167 non-duplicate samples (one re-sequenced sample failed QC and one original was missing) were flagged as potential swaps (5.4%), all originating from the same plate—suggesting plate-specific handling errors. In RS, 15 mismatches were identified among 127 samples (11.8%), with eight traced to a single plate. Notably, five of these re-sequenced RS samples were most similar to adjacent samples on the same plate, indicating possible plating shifts during pipetting or cross-contamination from neighboring wells.

Based on these findings, plates enriched for mismatches were prioritized for re-sequencing (i.e., second round of re-sequencing). Additionally, specific rows or columns within plate layouts that showed patterns of mismatched samples were also flagged and included in the re-sequencing list to further investigate and resolve potential systematic errors. Based on these findings, plates enriched for mismatches were prioritized for a second round of re-sequencing. Specific rows or columns within plate layouts showing patterns of mismatches were also flagged for further investigation. In GenR, 94 samples were re-sequenced, including 88 from the plate with nine mismatches. Although the initial plan was to resequence the entire plate (92 samples) four could not be processed due to limited DNA availability. Among these 88, 21 (23.8%) did not match their original counterparts, with most mismatches involving adjacent samples and concentrated in three columns of the plate. Six of these mismatches had corresponding laboratory annotations in the registry from the previous data release, suggesting procedural issues. Moreover in nine samples of this plate, where the original sample was missing (n=12), the replicate showed outlying proximity to another sample, further supporting the presence of swaps. Of the remaining six samples scattered in other plates, one was confirmed as a swap. Overall, only one out of 14 previously re-sequenced samples (pilot study) failed to match its earlier replicate.

In RS, two full plates and two columns from additional arrays were selected for re-sequencing. Due to insufficient DNA in some cases, random samples from other plates were included. Ultimately, 183 samples were incorporated into the microbiome swap identification pipeline. Thirty-five of these could not be compared to their original counterpart because the originals failed QC (see previous section). Among these, only three showed outlying proximity to other samples. Of the remaining 148 samples, 18 (12.2%) were flagged as potential mismatches, with 10 having laboratory annotations. In 13 additional cases, visualization of the similarity matrix revealed outlying proximity to other samples in the dataset.

A third re-sequencing wave was conducted, with the goal of maximizing the number of usable samples for downstream analyses. This effort focused on samples that had previously failed QC and those with unclear mismatch origins—likely due to pipetting errors in either of the earlier sequencing rounds. In total, 300 samples from Generation R were included in this third wave, from which 290 passed QC filters. Of these, 150 could not be compared to their original counterparts because the originals had failed sequencing, did not reach the minimum threshold of 4,000 reads, or were excluded based on other QC criteria. Despite this, eight of these re-sequenced samples (5.3%) showed outlying proximity to another sample in the dataset, suggesting potential swaps. Among the remaining 140 samples with available original counterparts, 11 (7.8%) were flagged as mismatches. Additionally, eight samples that had been previously re-sequenced showed high similarity to their earlier replicates, supporting their validity.

Altogether, we excluded samples confirmed as swaps, also those whose re-sequenced profiles did not match across different sequencing waves, those for which the original sample was missing but showed outlying proximity to another sample in the dataset, and those that failed the quality control filters described earlier. Additionally, samples which were identified as duplicates, with annotations from the collection center or study data management indicating procedural concerns, incomplete deposition records, or prolonged TimeInMail —defined as more than 7 days for RS and more than 11 days for GenR—were also excluded from the final dataset. After implementing all QC steps, GenR dataset comprised data derived from 2,526 participants (415 samples more than the previous release) while the RS dataset comprised profiling of 1,420 participants (7 samples less than the initial release). A summary of all the QC steps is shown in **Table 1**.

**Table 1.** Overview of result derived from each QC step.

|  | **Generation R Study** | **Rotterdam Study** |
| --- | --- | --- |
| Sample size of original datasets | 2,921 | 1,691 |
| Samples with successful sex determination assay | 2,851 (97.6%) | 1,445 (85.5%) |
| Sex-mismatches | 81 (2.8%) | 72 (5.0%) |
| Number of randomly selected samples for first round re-sequencing<sup>1</sup> | 350 | 350 |
| - Number of remaining samples after QC and exclusion of duplicates | 168 | 127 |
| -Number of samples where the original is in the dataset | 167 | 127 |
| - Number of detected sample swaps by beta diversity distance comparison | 9 (5.4%) | 15 (11.8%) |
| Number of selected samples for the second-round re-sequencing | 100 | 221 |
| -Number of remaining samples after QC | 94 | 183 |
| -Number of samples where the original is in the dataset | 80 | 148 |
| -Number of detected sample swaps by beta diversity distance comparison | 22 (27.5%) | 18 (12.2) |
| Number of selected samples for the third-round re-sequencing | 300 | 300 |
| -Number of remaining samples after filtering at 4K reads | 290 | 257 |
| -Number of samples where the original is in the dataset | 140 | 184 |
| -Number of detected sample swaps by beta diversity distance comparison | 11 (7.8%) | 40 (21.7%) |
| Number of recovered low-quality samples (reads <4K) from all rounds of re-sequencing | 343 | 317 |
| Total number of removed sample swaps<sup>3</sup> and low-quality samples after all QC steps | 738 | 588 |
| Sample size of final datasets after all QC steps | 2,526 | 1,420 |

“1”: The number has excluded samples that failed the sequencing during re-sequencing procedure.
“2”: The number has included 15 pairs of duplicates in GenR, and 7 pairs of duplicates in RS which are for algorithm validation purpose.
“3”: The sum of sample swaps included both swaps detected by sex-test assay and beta-diversity metrics.

### 3.3 Generation R and Rotterdam Study final datasets

A description of the generated datasets including sample size, number of taxonomies, and alpha diversity can be found in **Table 2**. In this new release, a total number of 1,578 ASVs and 1,759 ASVs were identified in the GenR and RS cohorts, respectively. Of these, 1,043 ASVs were shared across both cohorts, with 535 ASVs unique to GenR and 716 unique to RS. in the past year, probiotic use in the past three days, and key technical covariates—are summarized in **Table 3**.

**Table 2.** Overview of quality metrics for the updated microbiome datasets from the Generation R Study (GenR) and the Rotterdam Study (RS).

|  | GenR | RS |
| --- | --- | --- |
| Sample size | 2,526 | 1,420 |
| <sup>1</sup>Number of taxonomies | 1,578 | 1,759 |
| <sup>2</sup>Shannon index (mean ± sd) | 3.97 ± 0.42 | 3.97 ± 0.44 |
| <sup>2</sup>Richness (mean ± sd) | 146.21 ± 47.94 | 156.68 ± 55.56 |

“1”: number of amplicon sequence variants (ASVs).
“2”: Shannon index and richness were calculated at ASV level without rarefaction.

**Table 3.**
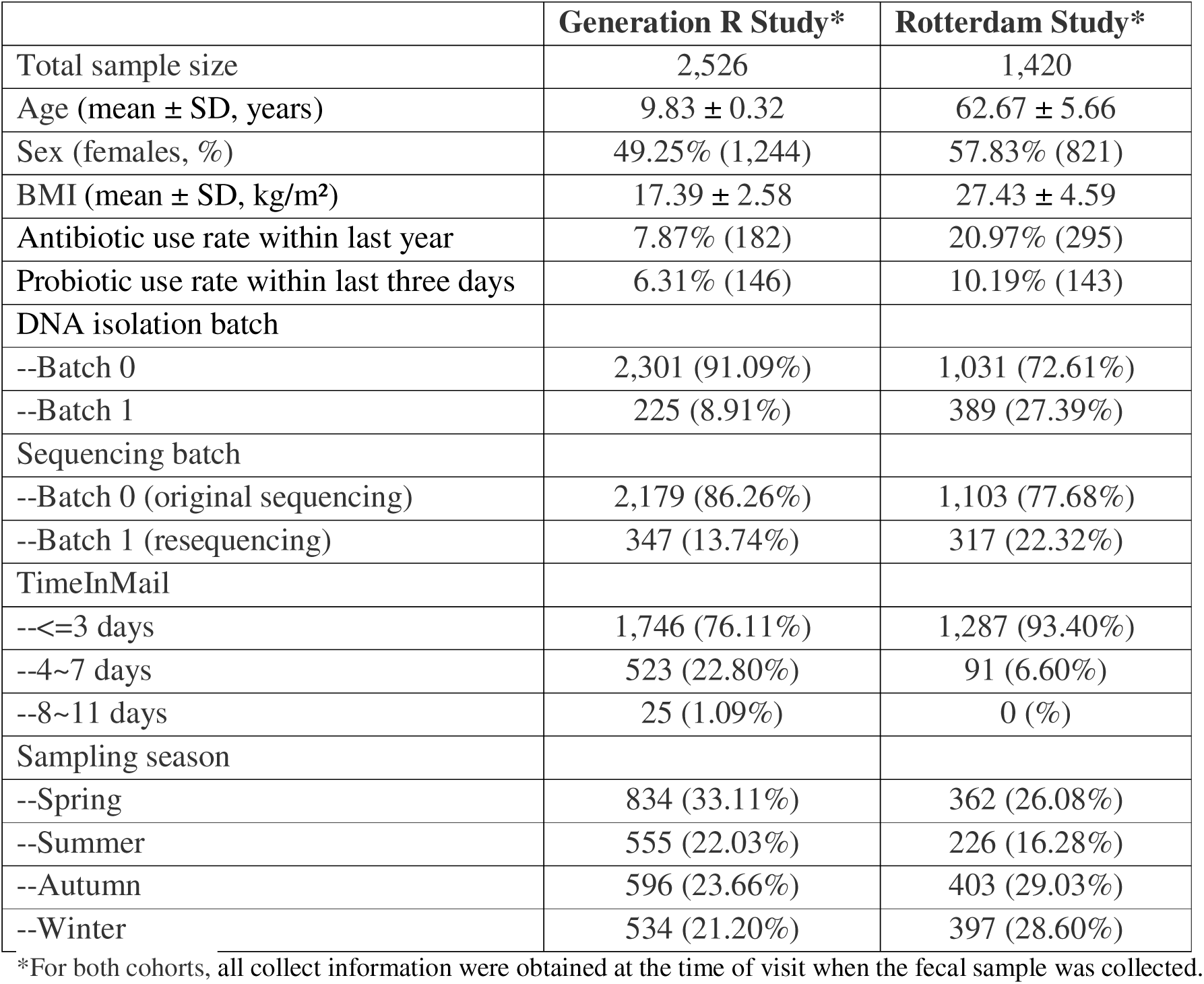
Basic characteristics of participants included in the updated microbiome datasets from the Generation R Study and the Rotterdam Study.

A summary of the taxa composition of participants included in this release is presented in **Figure 3**. The five most abundant phyla in both GenR and RS were Firmicutes, Bacteroidetes, Proteobacteria, Actinobacteria, and Verrucomicrobia. Compared to RS, GenR exhibited a lower relative abundance of Firmicutes and a higher relative abundance of Bacteroidetes.

**Figure 3.**
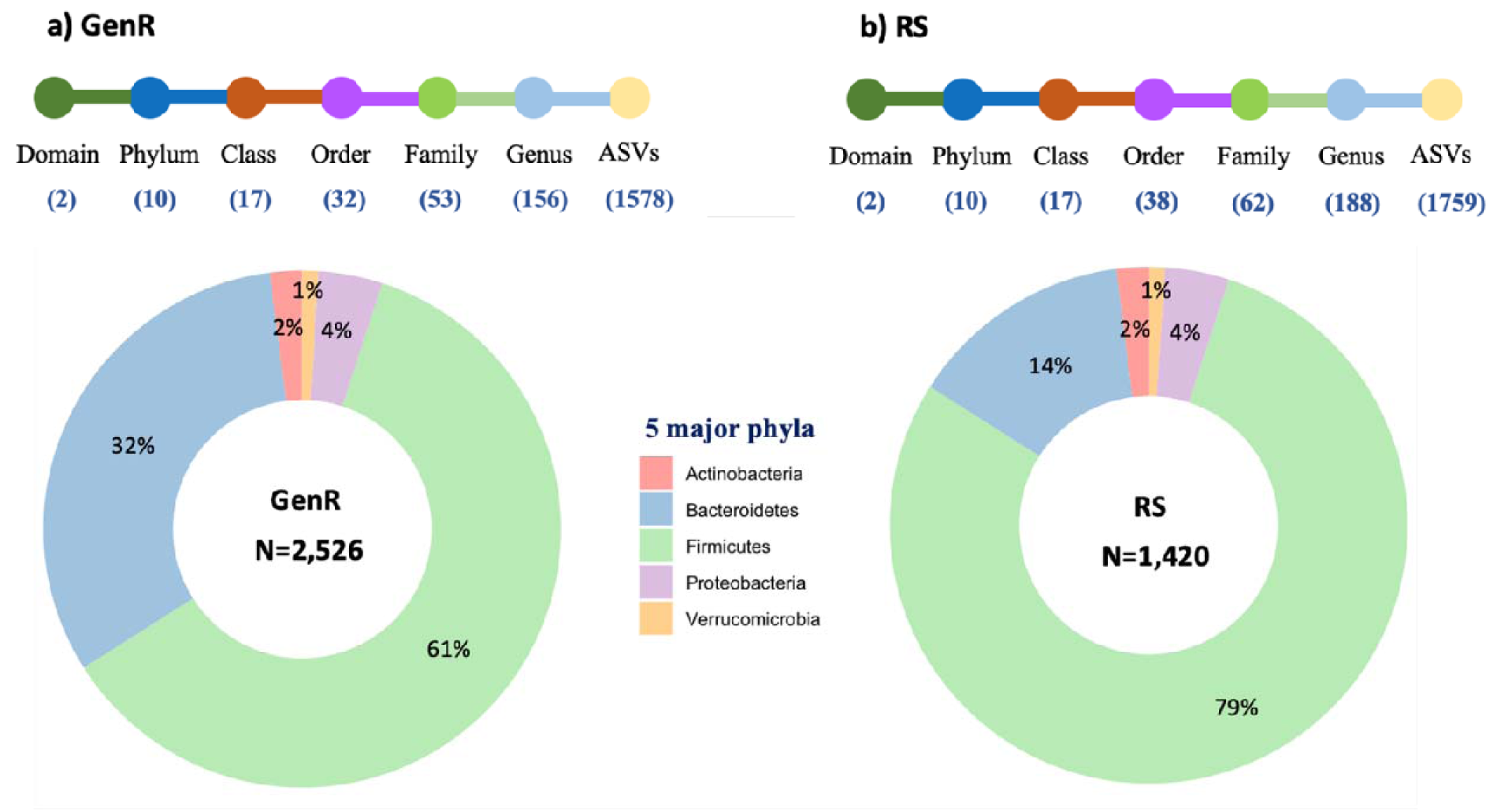
Overview of the most abundant Phyla in the Generation R Study and the Rotterdam Study. Top: number of observed taxa at each taxonomy level for each cohort. **Bottom:** Donut plots depicting the average relative abundances of the five major phyla in each cohort.

### 3.4 Validation of final 16S rRNA datasets of the GenR and RS cohorts

To validate the final datasets, we quantified the proportion of microbiome variance explained by technical covariates and individual phenotypes using PERMANOVA analyses (Figure 4). **Technical factors.** In the Generation R cohort, univariate analyses showed that DNA isolation batch, sequencing batch, number of reads, TimeInMail, and sampling season explained 0.80%, 0.35%, 0.38%, 0.26%, and 0.25% of the variance, respectively, with all technical covariates fitted together collectively accounting for 1.92% of the total microbiome variance. In the Rotterdam Study, DNA isolation batch, sequencing batch, number of reads, TimeInMail, and sampling season explained 0.23%, 1.00%, 0.35%, 0.27%, and 0.47% of the variance, respectively, totaling 2.2% when fitted collectively. Although different technical variables were evaluated in LL, comparable factors showed similar variance explained: batch (0.44%), number of reads (0.10%), and sampling season (0.36%). When including additional variables not assessed in our cohorts (DNA concentration, stool Bristol characterization, and deposition frequency), technical factors in LL collectively explained 2.32% of the total microbiome variance—comparable to our findings.

**Figure 4.**
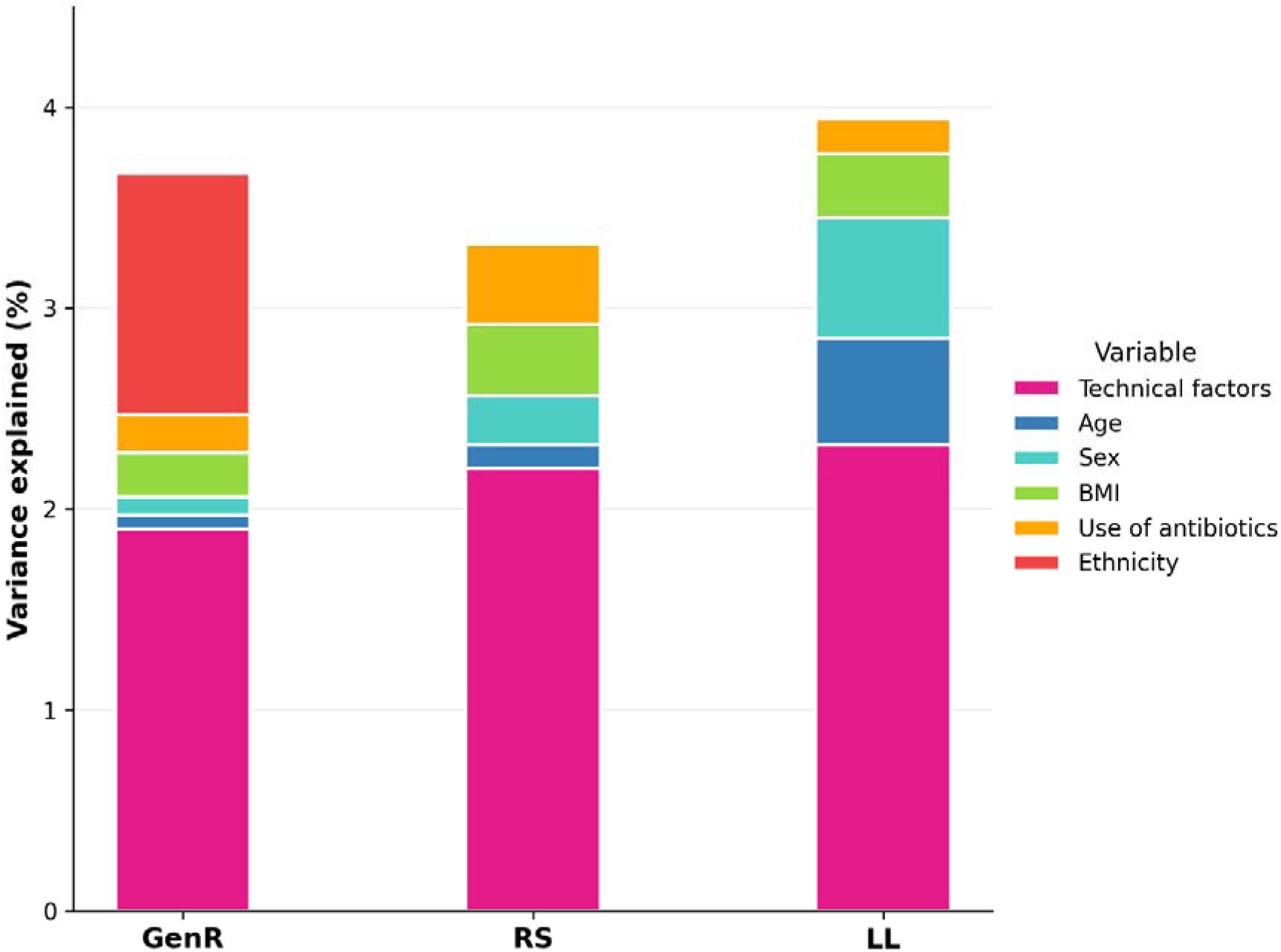
Variance in microbiome composition explained by technical factors and selected phenotypes. Bars represent variance explained by technical factors collectively and by individual phenotypes based on univariate PERMANOVA analysis. For RS and GenR, technical factors included DNA isolation batch, sequencing batch, number of reads, TimeInMail, and sampling season. For LL, technical factors included DNA isolation batch, number of reads, and sampling season.

**Individual phenotypes.** In GenR, age, sex, BMI and antibiotic use explained 0.07%, 0.09%, 0.22%, and 0.19% of the beta-diversity variance, respectively. In RS, these same factors explained 0.12%, 0.24%, 0.36%, and 0.40%, respectively. In LL, the corresponding values were 0.53%, 0.60%, 0.32%, and 0.17%. As Generation R is the only multi-ethnic cohort among the three, we additionally found that ethnicity explained 1.20% of the microbiome variance in this cohort.

## 4. Discussion

We have released a new freeze of microbiome data for the Rotterdam Study (N=1,420 adults, 1,759 ASVs) and Generation R Study (N=2,526, 1,578 ASVs), incorporating an improved denoising pipeline for ASV generation and an updated taxonomic classification database. Generating high-throughput sequencing data for thousands of samples is a complex, multi-step process that introduces substantial risk of sample mix-ups. Our workflow addresses this challenge through two complementary quality control strategies that detect and resolve sample swaps—a common yet under-addressed limitation in large population-based sequencing datasets. We also recovered some low-quality samples through re-sequencing, though samples with insufficient DNA available were missed.

Despite rigorous quality assurance measures, sample misidentification remains a challenge in large biobanks, and error rates are likely to increase as biobanking expands. Undetected sample swaps can lead to erroneous conclusions across multiple publications, making dataset validation critical for epidemiological and translational research. In this study, we focused on identifying and resolving sample swaps in two large microbiome cohorts. Our pilot study revealed a potential swap rate of 5.6-10%—higher than the ∼2% reported in transcriptomic studies using sex-discordance checks [43] or the ∼3% in genomic datasets [44]. This elevated rate likely reflects our study’s logistical complexity: home-based sample collection by thousands of participants, postal delivery and sorting at a collection center, before arrival to the sequencing facility for processing, therefore creating multiple handling steps where mislabeling could occur.

Our first quality control strategy leveraged host DNA traces from intestinal epithelial cells present in fecal samples. Although host DNA amounts vary by individual and health status [45], we successfully used this material to determine sex in most cases, enabling detection of sex-swaps. While the low coverage of host DNA in our cohorts precluded genome-wide variant calling, previous studies have demonstrated that human reads in metagenomic datasets can support SNP validation against blood-derived genotypes in both [45–47] both in low-biomass (cheeks and nostrils) [47] and gut microbiome samples [45, 46]. To our knowledge, this is the first study to systematically mine host genetic material from 16S rRNA-extracted DNA for participant re-identification as a quality control measure. Importantly, while host DNA serves as a valuable QC tool, it also poses privacy risks, underscoring the need for reliable computational removal in shared metagenomic datasets [48].

Because sex-based checks detect only opposite-sex swaps, we implemented a second strategy using beta-diversity dissimilarity matrices. This approach exploits the principle that biological replicates cluster more closely together than with non-replicates, enabling detection of all swap types regardless of sex. We validated this method by introducing duplicate samples in our pilot study, comparing sequencing profiles across samples with ≥4,000 reads after rarefying to equal depth. Although resequencing every sample in a large-scale study is cost-prohibitive, our pilot analysis of <25% f samples reliably estimated cohort-wide swap rates.

An important limitation is that our pipeline detected only post-DNA-isolation swaps, as we started from DNA isolated and stored 7 years ago. Other swap types potentially remaining in our datasets include household mix-ups (e.g., siblings/partners sharing households), registration errors during laboratory data entry, or mix-ups during manual sample preparation for DNA isolation in 12-tube racks. preparation of samples for DNA isolation that back in the day was done manually in racks containing 12 lose tubes. Obviously, the likelihood of such non-traceable swaps increases with the increase in size of the biobanks. Conversely, swaps during DNA dilution, pipetting, and templating are traceable with our approach. The likelihood of non-traceable swaps increases with biobank size, emphasizing the need for prevention through stringent laboratory procedures, workflow automation, and meticulous record-keeping throughout the sample lifecycle. Laboratory workflows, including ours, have improved substantially since data generation, likely reducing swap rates in current collections.

Beyond quality control, we evaluated technical and biological factors influencing microbiome profiles. Despite the low proportion of microbiome variance explained by the technical covariates surveyed here, given the complex nature of the dynamic microbiome data, we suggest researchers interested in the generated microbiome datasets to adjust for the DNA isolation batch, sequencing batch, time in mail, and sampling season to minimize unwanted variation and maximize power for detecting associations.

These datasets were intentionally filtered using lenient criteria to serve as a flexible resource for diverse research questions. Our thresholds intend to maximize retention of potentially biologically relevant low-abundance taxa while removing spurious ASVs. Common tools used for microbiome analysis such as ANCOM-BC or DESeq2, can handle sparcity well while might also benefit from more comprehensive data t calculate metrics such as the sampling fraction or size factor estimator intended to correct for library size variation or differential sampling efficiency. We recommend analysts to apply additional filtering appropriate to their specific research question. For association analyses, we recommend prevalence thresholds of 5-10% to balance statistical power with multiple testing burden

In summary, we applied updated pipelines and stringent QC procedures to release improved microbiome datasets for RS and GenR. Our scalable framework for identifying and resolving sample swaps is readily implementable in most laboratories and represents a practical approach to enhancing data quality in large population-based microbiome studies.

## CRediT authorship contribution statement

R.L. conceptualized the research idea, did the data analysis, and wrote the draft; A.K., A.H. and S.Y did the part of the data analysis; F.A., G.R., L.D. and K.A. reviewed and revised the manuscript; R.K., M.K, and A.U, reviewed the manuscript, provided supervision; V.J., J.F, and A.Z. reviewed the manuscript. C.M-G verified the data in this draft, reviewed and edited the draft, and provided supervision; F.R. reviewed and edited the draft, provided supervision and administrated this project.

## Declaration of Competing Interest

All authors declare that they have no competing financial interests or personal relationships that could have appeared to influence the work reported in this paper.

***All authors confirm they had full access to all the data in the study and accept responsibility to submit for publication*.**

## Acknowledgement

The generation and management of the 16S microbiome data for the Generation R Study was executed by the Human Genotyping Facility of the Genetic Laboratory of the Department of Internal Medicine, Erasmus MC, University Medical Center Rotterdam, the Netherlands. We thank Nahid El Faquir and Jolande Verkroost-Van Heemst for their help in sample collection and registration and Kamal Arabe, Hedayat Razawy, Karan Singh Asra, Pelle van der Wal, Sergio Chavez Chavez and Djawad Radjabzadeh for their help in DNA isolation and sequencing, and Joost Verlouw, Dr. Constanza Vallerga and Marijn Verkerk for their help with the bioinformatic analyses. We thank Ruolin Li, Dr. Cindy Boer, Dr. Robert Kraaij, Dr. Carolina Medina-Gomez and Prof. Joyce van Meurs for overseeing the quality control of the generated datasets. Furthermore, we express our gratitude to Drs Jeroen Raes and Jun Wang (Katholieke Universiteit Leuven, Belgium) for their guidance in 16S rRNA profiling and dataset generation. The Generation R Study gratefully acknowledge the contribution of children, parents and midwives, and general practitioners, hospitals, and pharmacies in Rotterdam. Also, the Rotterdam Study gratefully acknowledge the contribution of the inhabitants, general practitioners, and pharmacists of the Ommoord district.

The general design of the Generation R Study was made possible by financial support from the Erasmus MC, Rotterdam, the Erasmus University Rotterdam, the Netherlands Organization for Health Research and Development (ZonMW), the Netherlands Organization for Scientific Research (NWO), the Ministry of Health, Welfare and Sport and the Ministry of Youth and Families. The Rotterdam Study is funded by Erasmus Medical Center and Erasmus University, Rotterdam, Netherlands Organization for the Health Research and Development (ZonMw), the Research Institute for Diseases in the Elderly (RIDE), the Ministry of Education, Culture and Science, the Ministry for Health, Welfare and Sports, the European Commission (DG XII), and the Municipality of Rotterdam.

## Declaration of Competing Interest

All authors confirm they had full access to all the data in the study and accept responsibility to submit for publication.

## Acknowledgement

The generation and management of the 16S microbiome data for the Generation R Study and the Rotterdam Study was executed by the Human Genotyping Facility of the Genetic Laboratory of the Department of Internal Medicine, Erasmus MC, University Medical Center Rotterdam, the Netherlands. We thank Nahid El Faquir and Jolande Verkroost-Van Heemst for their help in sample collection and registration and Kamal Arabe, Hedayat Razawy, Karan Singh Asra, Pelle van der Wal, Sergio Chavez Chavez and Djawad Radjabzadeh for their help in DNA isolation and sequencing, and Dr. Constanza Vallerga and Marijn Verkerk for their help with the bioinformatic analyses. The Generation R Study gratefully acknowledge the contribution of children, parents and midwives, and general practitioners, hospitals, and pharmacies in Rotterdam. Also, the Rotterdam Study gratefully acknowledge the contribution of the inhabitants, general practitioners, and pharmacists of the Ommoord district.

## Data availability

Because of restrictions based on privacy regulations and informed consent of the participants, data cannot be made freely available in a public repository. However, data from this study can be obtained upon reasonable request to the director of the Generation R Study, subject to local, national and European rules and regulations.

